# Expansion of a fly TBI model to four levels of injury severity reveals synergistic effects of repetitive injury for moderate injury conditions

**DOI:** 10.1101/611244

**Authors:** Lauren J Putnam, Ashley M Willes, Brooke E Kalata, Nathaniel D Disher, Douglas J Brusich

## Abstract

Several million traumatic brain injury (TBI) events are reported in the United States annually. However, mild TBI events often go unreported, and mild and repetitive mild TBI conditions are challenging to model. Fruit flies (*Drosophila melanogaster*) have gained traction for the study of TBI. The best-characterized fly TBI model is the high-impact trauma (HIT) method. We replicated the HIT method and confirmed several previous findings at the standard level of injury severity. We then expanded upon the HIT model by characterizing mortality across three reduced levels of injury severity. Importantly, we found reduced mortality with reduced injury severity and synergistic effects on mortality in response to repetitive TBI by our moderate injury conditions. Last, we compared moderate, repetitive TBI to a single severe TBI via assessment of the pattern of mortality and geotaxis performance in the 24 h following TBI. We found the number and severity of injuries could result in different patterns of death, while all TBI conditions led to impaired geotaxis compared to uninjured flies at 0.5 h and 6 h post-TBI. Thus, we have extended a well-characterized model of TBI in flies, and shown the utility of this model for making unique insights into TBI across various severities, injury numbers, and time-points post-injury.

## INTRODUCTION

In the United States, traumatic brain injury (TBI) annually accounts for greater than 2.5 million emergency room (ER) visits, hospitalizations, and deaths combined (Taylor et al. 2017). Additionally, half of all mild TBI events are estimated to go unreported (Cassidy et al. 2004). Mild TBI events, including concussion and sub-concussive impacts, are commonly suffered during sports participation and military deployment (Marar et al. 2012; Helmick et al. 2015; Kerr et al. 2017; Baldwin et al. 2018). Individuals in contact sports such as football may experience greater than one thousand mild head impacts per year, while approximately 10% of U.S. Army soldiers reported multiple mild TBI events from a previous deployment (Crisco et al. 2010; Wilk et al. 2012).

Severe TBI events are associated with long-term outcomes including greater risk for dementia, and are associated with many hallmarks of neurodegenerative disease (DeKosky and Asken 2017; Nordström and Nordström 2018). Individual mild TBI events are not well-linked to long-term outcomes, and most TBI-associated conditions resolve within months, particularly in children (Holm et al. 2005). By contrast, repetitive mild TBI is associated with more prominent impairment or disease, such as a greater risk of neurodegenerative disease in American football players (Lehman et al. 2012; Bailes et al. 2013; Levin and Robertson 2013). Moreover, rodent models of repetitive mild TBI result in neurocognitive deficits, and histological and morphological changes associated with neurodegenerative disease (Mouzon et al. 2012; Ojo et al. 2016; Gold et al. 2018). Importantly, additional TBI events suffered within days of the first injury result in more negative outcomes due to the combination of primary and secondary injury mechanisms (Laurer et al. 2001; Longhi et al. 2005; Friess et al. 2009; Meehan et al. 2012; Huang et al. 2013; Bolton and Saatman 2014; Weil et al. 2014; Bolton Hall et al. 2016).

Two models for studying TBI have been developed for the fruit fly (*Drosophila melanogaster*): the high-impact trauma (HIT) method, which uses a spring-based device deflected to 90°, and the Bead Ruptor homogenizer method, which uses a programmable homogenizer that can be set to various speeds and durations (Katzenberger et al. 2013; Barekat et al. 2016). Importantly, use of each method results in classic post-TBI symptoms including impaired locomotion, shortened lifespan, neurodegeneration, intestinal barrier disruption, and activation of immune and autophagy processes (Katzenberger et al. 2013; Barekat et al. 2016; Anderson et al. 2018). While the Bead Ruptor method offers potential advantages in the ease of scaling primary injuries and inter-experiment standardization, the HIT method is simple, cost-effective, and better characterized to date (Katzenberger et al. 2013, 2015, 2016; Barekat et al. 2016; Anderson et al. 2018).

We sought to standardize the HIT method across several levels of injury severity, thereby extending this well-characterized method to the study of mild to severe TBI events in an easily replicable manner. To this end, we installed fixed, selectable stopping points that limited deflection of the HIT device to either 60°, 70°, 80°, or 90°. We found that reducing the angle of deflection greatly reduced mortality, and that repetitive injury at moderate levels of severity resulted in a pronounced synergistic effect on mortality. Moreover, we found that the pattern of death in the 24 h post-TBI could be affected by the nature of repetitive injury. Last, we found that locomotion was impaired when assessed at an early 0.5 h time-point and also during the secondary injury window at 6 h post-TBI, but was no different than controls when assessed 2 h or 24 h post-TBI.

## MATERIALS AND METHODS

### Fly Husbandry

Flies of genotype *w*^*1118*^ (BL 5905) and *y*^*1*^*w*^*1*^ (BL 1495) were obtained from the Bloomington Drosophila Stock Center (Bloomington, Indiana, USA). Flies were maintained in a 25°C humidified incubator on a 12H:12H light:dark cycle. Flies were maintained on a glucose-cornmeal-yeast media with the following quantities per 1.25L of water: 7.66g agar (Apex), 14.4g glucose (DOT Scientific Inc.), 50.3g cornmeal (Genesee), 15g yeast (Genesee), 5.66mL tegosept (Genesee), 4.67mL propionic acid (99%, Acros Organics), and 0.47mL phosphoric acid (85%, Matheson Coleman & Bell Inc.).

### TBI Methodology

Flies were collected using light-CO_2_ anesthesia. Flies were subjected to traumatic brain injury on or before 5 days after eclosion (dae) using an adapted model of the high-impact trauma (HIT) device (Katzenberger et al. 2013). Briefly, flies were transferred to an empty vial and the vial was affixed to the end of a compression spring. The vial was deflected to a selectable, fixed stopping point of 60°, 70°, 80° or 90°. The vial was released and allowed to collide with a foam pad covered by a 1/16” rubber pad. Vial deflections were repeated every 15 seconds for the total number of deflections indicated. Flies were immediately hand-transferred to a food vial following the final injury. Uninjured flies were handled identically minus spring deflection and injury. The number of dead flies were counted at designated time-points ranging from 30 minutes post-injury to 24 h post-injury. Flies counted at 24 h post-injury were used to determine the mortality index at 24 h (MI_24_) (MI_24_ = # flies dead at 24 h post-injury/total # flies * 100). The MI_24_/HIT values were determined using the MI_24_ divided by total number of injuries for the condition.

### Negative Geotaxis

Flies aged 0-4 dae were collected under light-CO_2_ anesthesia and transferred to food vials. Animals were subjected to TBI, following the same methods as above, and geotaxis testing, at a designated time post-TBI, a minimum of 1 day after CO_2_ exposure. Individual vials of flies were tested at a single time-point each and a minimum of 5 vials were used for each time-point. Prior to geotaxis testing, flies were hand-transferred to empty vials and given 10 minutes undisturbed on the geotaxis platform to recover prior to experimentation. Vial plugs were kept to within the top 10mm of the vial.

A custom-built geotaxis apparatus which accommodated up to 6 vials simultaneously was used for all geotaxis testing. Briefly, the device consisted of wood and plywood construction affixed to two ring stands used as vertical runners. Geotaxis behavior was initiated by startle whereby the device was lifted to stops on the ring stands which allowed approximately 95mm of vertical movement, dropped, and the process repeated twice more for a total of 3 drops in quick succession. Foam pads were used to cushion both the device and the platform upon which the vials rested. Vials were held in place on the device by use of elastic cords placed around the top 10mm of the vial. A webcam (Logitech c270) was used to record each experiment.

VLC media player (version 3.0.6) with a self-written subtitle file displaying the video time to the tenths of seconds was used for analysis. Screenshots were taken at 5.0 s and 10.0 s after the 3^rd^ drop. Screenshots were processed in ImageJ (1.47v/Java 1.6.0_20 (32-bit)). Three lines were drawn to measure the height of the vial in pixels and averaged. The Cell Counter plug-in was used to mark vial bottoms and flies for each sample. Pixel coordinates from Cell Counter were combined with the pixel measure of the vial and the known length of the vial (95mm) to convert each fly’s pixel coordinate to a distance traveled from the vial bottom in millimeters. Distance measurements were capped at 50mm. GraphPad Prism 7 software (GraphPad Software, Inc.) was used to generate a histogram of distance measures and bin values into 12.5mm quartiles. Dead flies were not included in geotaxis measurements.

### Statistics

Pairwise comparisons of categorical (dead:alive) count data were performed using a 2×2 Fisher’s Exact Test between selected conditions (GraphPad Prism 7). Bonferroni correction was used to correct for multiple testing and corrected alpha levels are reported in figure legends. Comparisons of median MI_24_/HIT values and geotaxis data were conducted via Kruskal-Wallis testing (GraphPad Prism 7) with multiple comparisons of mean ranks and Dunn’s correction at a level of α = 0.05. Only vials containing at least 30 flies were used in median MI_24_/HIT comparisons. Full count data were used for comparisons of trends across MI_24_/HIT data; overall MI_24_ values were divided by their respective HIT number, plotted across 1-4HITs, and fitted using the linear fit mode within the nonlinear regression analysis toolkit (GraphPad Prism 7). Lines were fitted using the least squares fit mode, compared to a hypothetical slope of zero via the extra sum-of-squares F test at a level of α = 0.05, and the 95% confidence interval (CI) determined asymmetrically. Dead fly counts from determination of death across time-points were compared by Fisher’s Exact Tests of 4×2 matrices in R (version 3.5.1) with post-hoc, pairwise comparisons (dead within window:dead outside of window) via 2×2 Fisher’s Exact Tests (GraphPad Prism 7). Bonferroni correction was used to correct for multiple post-hoc testing and the corrected alpha level is reported in the text or figure legend.

## RESULTS

### Replication of 90° HIT data

The primary measure of TBI outcomes in flies is the percentage of flies that die within 24 hours post-injury (MI_24_). We first set out to determine how our TBI system and resulting MI_24_ values compared to existing models. We conducted experiments using a 90° angle of deflection and two strains of fruit fly, *w*^*1118*^ and *y*^*1*^*w*^*1*^, for which *y*^*1*^*w*^*1*^ was previously reported to suffer higher MI_24_ (Katzenberger et al. 2013). Vials of flies were subjected to 0-4 high-impact traumatic injuries (HITs) (see Table 1 for all categorical count data). Uninjured flies suffered little or no mortality at 24 h, while administration of 1-4HITs resulted in pronounced MI_24_ with increased death upon increased HIT number (Fig. 1A, shared letters indicate statistical significance between conditions). Comparisons across genotypes showed MI_24_ values of *w*^*1118*^ and *y*^*1*^*w*^*1*^ flies were no different for uninjured controls, but *y*^*1*^*w*^*1*^ flies suffered greater mortality than *w*^*1118*^ flies for each of the 1-4HIT datasets (Fig. 1A, (*) indicates differences between genotypes).

**Table 1:**
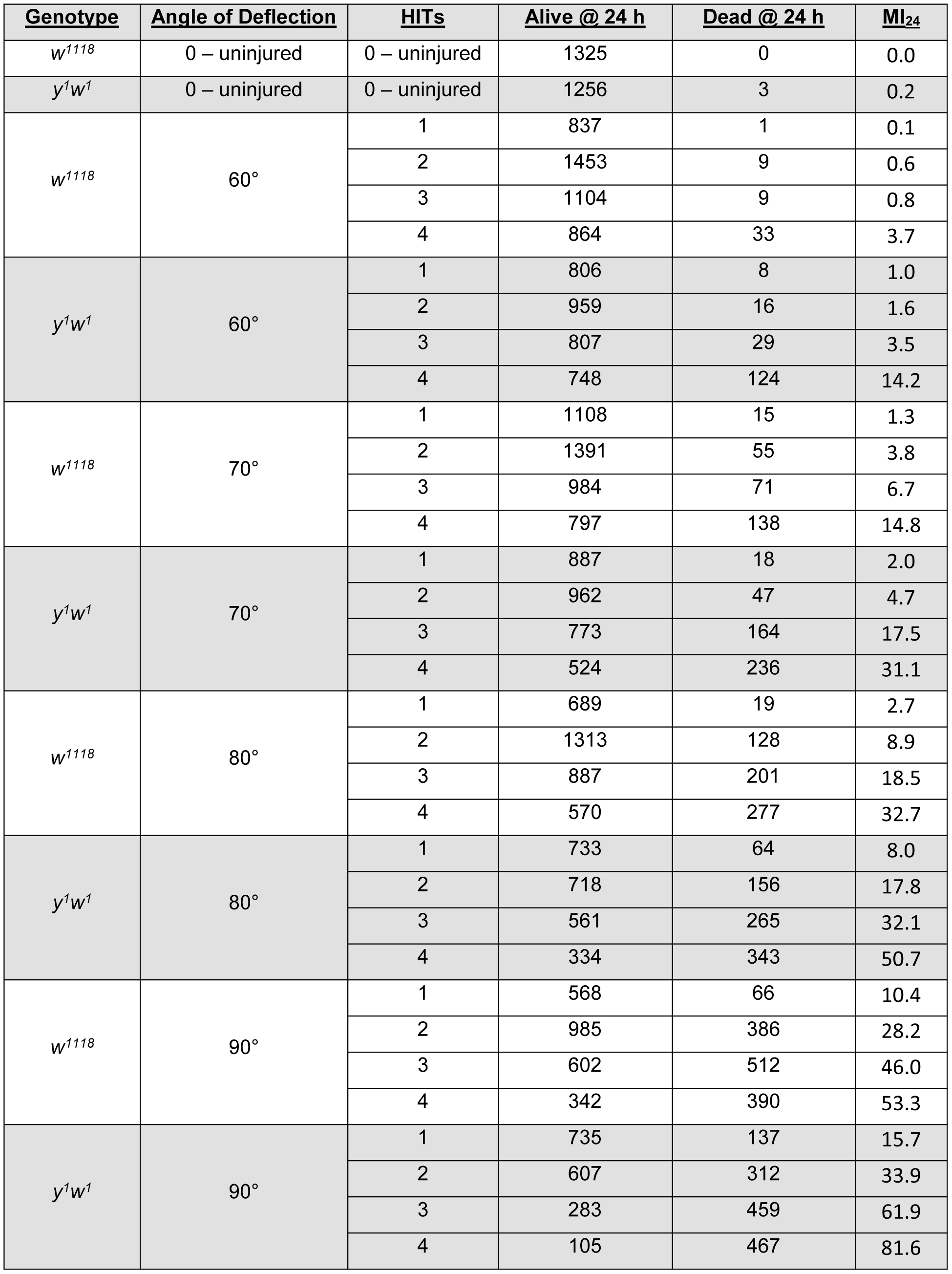
Full reporting of categorical count data for all TBI conditions.

| <u>Genotype</u> | <u>Angle of Deflection</u> | <u>HITs</u> | <u>Alive @ 24 h</u> | <u>Dead @ 24 h</u> | <u>MI<sub>24</sub></u> |
| --- | --- | --- | --- | --- | --- |
| $w^{1118}$ | 0 – uninjured | 0 – uninjured | 1325 | 0 | 0.0 |
| $y^1w^1$ | 0 – uninjured | 0 – uninjured | 1256 | 3 | 0.2 |
| $w^{1118}$ | 60° | 1 | 837 | 1 | 0.1 |
|  |  | 2 | 1453 | 9 | 0.6 |
|  |  | 3 | 1104 | 9 | 0.8 |
|  |  | 4 | 864 | 33 | 3.7 |
| $y^1w^1$ | 60° | 1 | 806 | 8 | 1.0 |
|  |  | 2 | 959 | 16 | 1.6 |
|  |  | 3 | 807 | 29 | 3.5 |
|  |  | 4 | 748 | 124 | 14.2 |
| $w^{1118}$ | 70° | 1 | 1108 | 15 | 1.3 |
|  |  | 2 | 1391 | 55 | 3.8 |
|  |  | 3 | 984 | 71 | 6.7 |
|  |  | 4 | 797 | 138 | 14.8 |
| $y^1w^1$ | 70° | 1 | 887 | 18 | 2.0 |
|  |  | 2 | 962 | 47 | 4.7 |
|  |  | 3 | 773 | 164 | 17.5 |
|  |  | 4 | 524 | 236 | 31.1 |
| $w^{1118}$ | 80° | 1 | 689 | 19 | 2.7 |
|  |  | 2 | 1313 | 128 | 8.9 |
|  |  | 3 | 887 | 201 | 18.5 |
|  |  | 4 | 570 | 277 | 32.7 |
| $y^1w^1$ | 80° | 1 | 733 | 64 | 8.0 |
|  |  | 2 | 718 | 156 | 17.8 |
|  |  | 3 | 561 | 265 | 32.1 |
|  |  | 4 | 334 | 343 | 50.7 |
| $w^{1118}$ | 90° | 1 | 568 | 66 | 10.4 |
|  |  | 2 | 985 | 386 | 28.2 |
|  |  | 3 | 602 | 512 | 46.0 |
|  |  | 4 | 342 | 390 | 53.3 |
| $y^1w^1$ | 90° | 1 | 735 | 137 | 15.7 |
|  |  | 2 | 607 | 312 | 33.9 |
|  |  | 3 | 283 | 459 | 61.9 |
|  |  | 4 | 105 | 467 | 81.6 |

**Figure 1:**
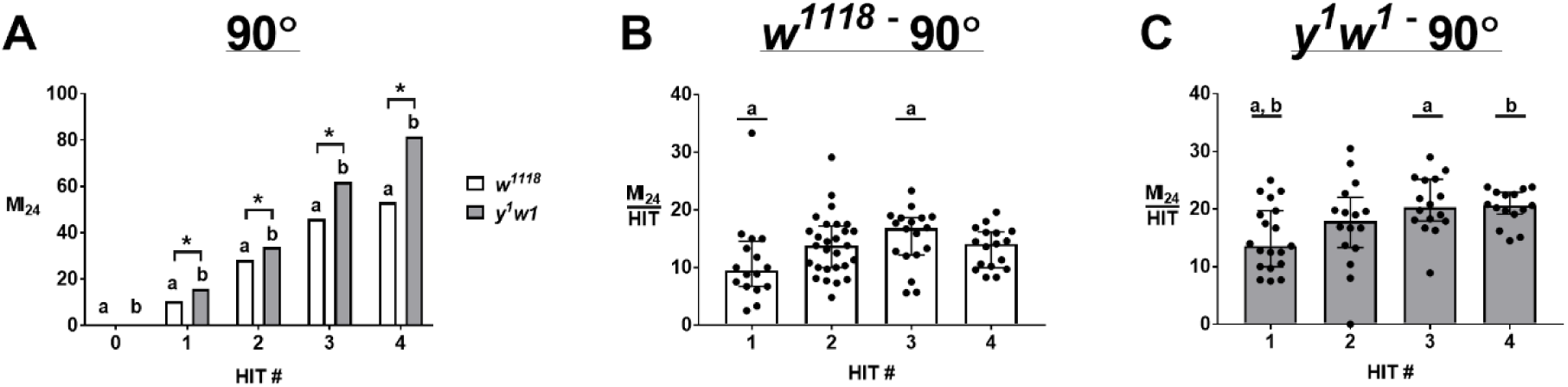
Increasing HIT number at 90° deflection increases MI_24_ and reveals differences in MI_24_ per hit. (A) MI_24_ values increase with HIT number in both *w*^*1118*^ and *y*^*1*^*w*^*1*^, with *y*^*1*^*w*^*1*^ suffering greater MI_24_ across all HIT numbers. Zero HITs represents uninjured controls. Conditions that share a letter are statistically different (p ≤ 0.0023, α = 0.005), while (*) indicates differences between genotypes (p ≤ 0.0029, α = 0.01) by Fisher’s Exact Test with Bonferroni correction. n ≥ 572 flies for each condition. (B, C) MI_24_ values were divided by HIT number for *w*^*1118*^ (B) and *y*^*1*^*w*^*1*^ (C). Data plotted are medians with interquartile ranges, with individual data points for each vial of at least 30 flies. Conditions that share a letter are statistically different (p < 0.05, α = 0.05 by Kruskal-Wallis with Dunn’s correction, n ≥ 15 vials for each condition).

It was previously reported that MI_24_ values divided by HIT number (MI_24_/HIT) were no different from one another when compared across a range of HITs (Katzenberger et al. 2013). We carried out the same comparisons for our datasets and found differences for median MI_24_/HIT values for *w*^*1118*^ (Fig. 1B, 1HIT vs 3HIT) and *y*^*1*^*w*^*1*^ (Fig. 1C, 1HIT vs 3HITs, and 1HIT vs 4HITs). The differences in MI_24_/HIT prompted us to look more closely at the pattern of change in mortality across HIT numbers. If mortality is directly proportional to the number of flies which experience a critical injury for each HIT then we would expect the MI_24_/HIT values compared across HIT numbers to have a zero slope. We used overall count data to determine MI_24_/HIT values and then fitted these points across 1-4 HITs with a linear best-fit model. At 90° we found that neither *w*^*1118*^ nor *y*^*1*^*w*^*1*^ had slopes that significantly deviated from zero (Table 2).

**Table 2:** Full reporting of line-fit slopes for MI_24_/HIT data and resulting p-values.

| <u>Genotype</u> | <u>Angle of Deflection</u> | <u>Slope +/- SE</u> | <u>95% CI of slope</u> | <u>Significantly non-zero slope?</u> | <u>p-value</u> |
| --- | --- | --- | --- | --- | --- |
| $w^{1118}$ | 60 | 0.24 +/- 0.1 | -0.18 to 0.66 | no | 0.136 |
| $y^1w^1$ | 60 | 0.81 +/- 0.42 | -1.00 to 2.61 | no | 0.195 |
| $w^{1118}$ | 70 | 0.74 +/- 0.17 | 0.02 to 1.46 | **yes** | **0.048** |
| $y^1w^1$ | 70 | 2.08 +/- 0.42 | 0.28 to 3.89 | **yes** | **0.038** |
| $w^{1118}$ | 80 | 1.82 +/- 0.05 | 1.61 to 2.03 | **yes** | **0.001** |
| $y^1w^1$ | 80 | 1.57 +/- 0.18 | 0.81 to 2.33 | **yes** | **0.013** |
| $w^{1118}$ | 90 | 1.00 +/- 0.90 | -2.87 to 4.86 | no | 0.382 |
| $y^1w^1$ | 90 | 1.77 +/- 0.50 | -0.37 to 3.92 | no | 0.071 |

### Expansion to three levels of reduced injury severity

In order to expand the range of primary injury severities by the HIT method, we added additional, fixed, selectable stopping points to reduce the angle of deflection to 80°, 70°, or 60°. We again assessed MI_24_ outcomes by independently administering 1-4HITs at each of the three new angles of deflection. We found lower MI_24_ values in each of the new deflection angles when compared to 90° within both *w*^*1118*^ and *y*^*1*^*w*^*1*^ datasets at each of 1-4HITs (Fig. 2A-D respectively). We also found significantly reduced MI_24_ with each reduction in deflection angle from 80° to 70° and then 70° to 60° at each of 2-4 HITs within both *w*^*1118*^ and *y*^*1*^*w*^*1*^ datasets (Figs. 2B-D), while genotype-specific differences across deflection angle were seen at 1HIT (Fig. 2A). Moreover, in both genotypes we found the MI_24_ from 1HIT at 60°, our most mild injury severity, was not significantly different than the MI_24_ in uninjured animals using a significance level of α = 0.005 after Bonferroni correction (Fig. 2A, p-values: *w*^*1118*^ = 0.39, *y*^*1*^*w*^*1*^ = 0.03). Last, we found *y*^*1*^*w*^*1*^ flies suffered greater mortality than *w*^*1118*^ flies at all deflection angles when 3 or 4 HITs were administered (Figs. 2C and 2D). However, differences between genotypes were only statistically different at the 80° and 90° deflection angles when injuries were limited to 1 or 2 HITs (Figs. 2A and 2B). Nonetheless, *y*^*1*^*w*^*1*^ flies appeared comparatively more sensitive to TBI at less severe primary injuries as the fold-difference in *y*^*1*^*w*^*1*^:*w*^*1118*^ MI_24_ values progressively decreased from 3.84-fold at 60° to 1.53-fold at 90° for 4HITs (Fig. 2D), a pattern similarly observed for other HIT numbers.

**Figure 2:**
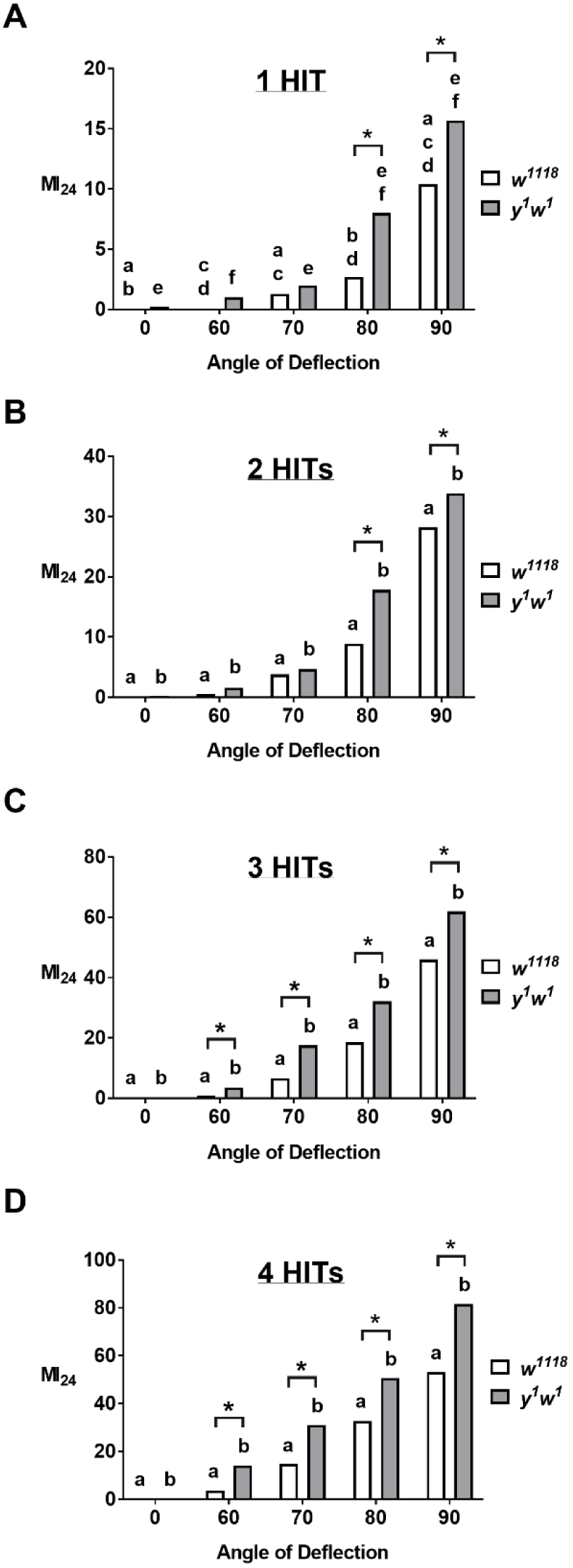
Mortality is reduced at smaller angles of deflection. Flies were administered 1-4HITs (A-D as indicated) at designated angles of deflection from 60° to 90°. Zero degrees represents uninjured controls. Conditions that share a letter are statistically different (p ≤ 0.0042, α = 0.005), while (*) indicates differences between genotypes (p ≤ 0.0035, α = 0.01) by Fisher’s Exact Test with Bonferroni correction. n ≥ 572 flies for each condition.

### Synergistic effects are apparent for repetitive injury at moderate TBI severity

We continued our analysis of the additional deflection angles to comparisons of MI_24_/HIT values. We found differences in MI_24_/HIT values when comparing 1HIT and 4HITs in both *w*^*1118*^ and *y*^*1*^*w*^*1*^ at each of the sub-90° deflection angles (Fig. 3). Additionally, differences between both 1HIT and 3HITs, and 2HITs and 4HITs were also seen for *w*^*1118*^ at 80° (Fig. 3A) and *y*^*1*^*w*^*1*^ at 70° (Fig. 3D). We again investigated trends in MI_24_/HIT data from 1-4HITs via analysis of slopes from best-fit lines. If mortality was strictly additive for each HIT then the trend across MI_24_/HIT data should generate a zero-slope line. However, at both 80° and 70°, but not 60°, both *w*^*1118*^ and *y*^*1*^*w*^*1*^ had positive, significantly non-zero slopes, indicating a synergistic effect on mortality (Table 2). The positive slopes and synergistic effects were most evident at moderate severity injuries of 80° for *w*^*1118*^ (1.82 +/- 0.05 (SE)) and 70° for *y*^*1*^*w*^*1*^ (2.08 +/- 0.42 (SE)) (Table 2).

**Figure 3:**
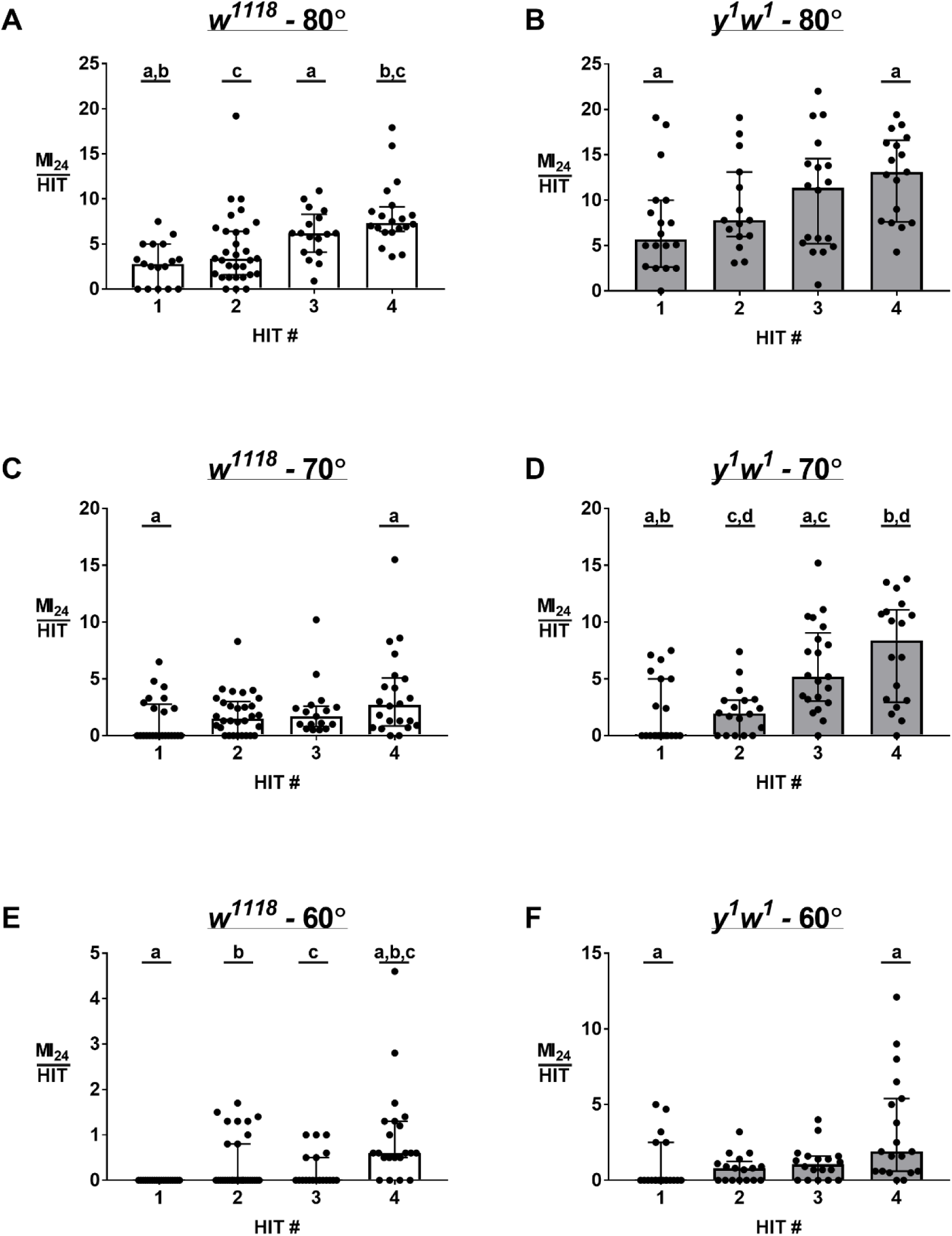
Differences in MI_24_ per hit are readily apparent for sub-90° injury conditions. MI_24_ values were divided by HIT number for *w*^*1118*^ and *y*^*1*^*w*^*1*^ as indicated at angles of deflection of 80° (A, B), 70° (C, D) and 60° (E,F). Data plotted are medians with interquartile ranges, with individual data points for each vial of at least 30 flies. Conditions that share a letter are statistically different (p < 0.05, α = 0.05 by Kruskal-Wallis with Dunn’s correction). n ≥ 15 vials for each condition.

### Time-Course of Mortality

Animals subjected to TBI experience both primary injury from the TBI event itself and secondary injuries related to cellular and molecular events instigated by the primary injury. The secondary injury period in flies reportedly peaks between 1 h and 8 h post-injury and persists for at least 24 h (Katzenberger et al. 2016). We hypothesized that the relative contributions of primary and secondary injuries on mortality would differ when comparing a single severe injury (90° × 1HIT) to repetitive injury at less severe angles. By our data, injury via 90° × 1HIT results in similar MI_24_ values as 80° × 2HITs and 70° × 3HITs, particularly for *y*^*1*^*w*^*1*^ flies (Table 1), thereby giving us conditions by which to test our hypothesis while keeping overall MI_24_ values comparable. We injured new cohorts of flies by these 3 conditions and recorded mortality post-TBI at times which corresponded to an early, pre-secondary injury window (0.5 h), early peak of the secondary injury period (2 h), delayed peak of the secondary injury period (8 h), and late secondary injury period (24 h).

We found the only conditions which generated different patterns in the time-course of death were 80° × 2HITs vs 90° × 1HIT, and this was true for both the *w*^*1118*^ and *y*^*1*^*w*^*1*^ genotypes (Fig. 4). Notably, while the overall trends between the two conditions differed for *w*^*1118*^ there were no pairwise differences by post-hoc testing (Fig. 4). By contrast, *y*^*1*^*w*^*1*^ flies showed significantly greater late death from 8 h – 24 h when injured by 80° × 2HITs vs. 90° × 1HIT (Fig. 4). Injury via 70° × 3HITs resulted in no differences in the time-course of death compared to either the 80° × 2HITs or 90° × 1HIT conditions for either genotype (Fig. 4).

**Figure 4:**
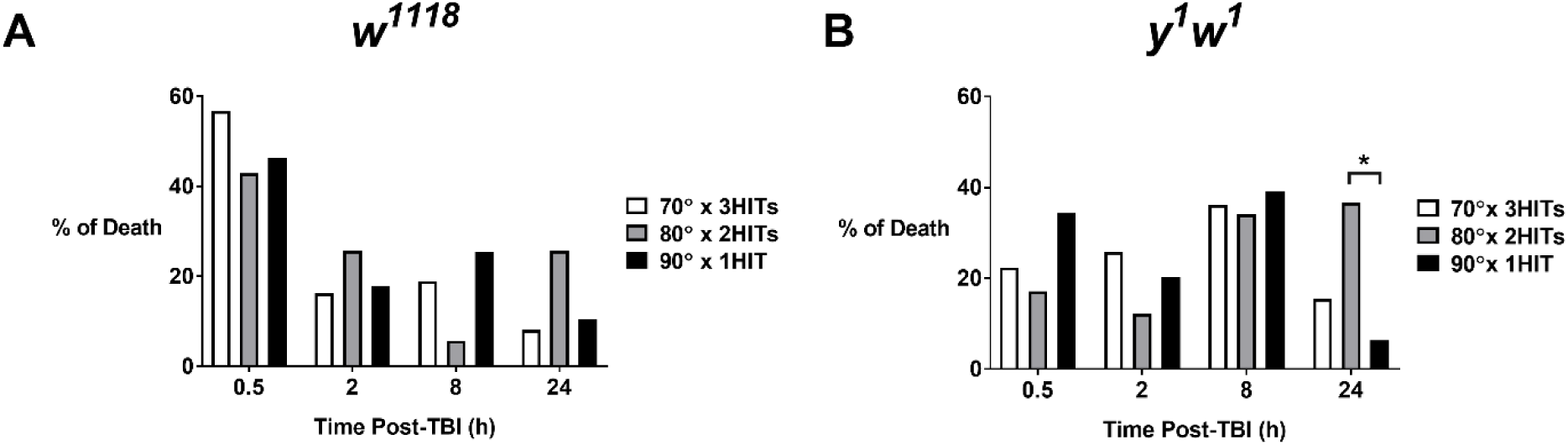
The time-course of mortality differs for 80° × 2HITs vs 90° × 1HIT. *w*^*1118*^ (A) and *y*^*1*^*w*^*1*^ (B) flies were injured by one of three designated conditions and the percentage of total death by 24 h post-injury is plotted for each of 4 time-points. Datasets of 80° × 2HITs vs 90° × 1HIT differed for each genotype (p = 0.026 for *w*^*1118*^, p = 0.001 for *y*^*1*^*w*^*1*^, α = 0.05 by 4×2 Fisher’s Exact Test), while (*) indicates a difference via post-hoc pairwise comparison (p < 0.001, α = 0.0125 by Fisher’s Exact Test with Bonferroni correction). n ≥ 276 flies per condition.

Last, we compared the pattern of death between the *w*^*1118*^ and *y*^*1*^*w*^*1*^ genotypes. We found the pattern of death from both the 70° × 3HITs and 80° × 2HITs differed between genotypes (Figs. 4A and 4B; 70°: p = 0.010, 80°: p = 0.002, α = 0.017 by 4×2 Fisher’s Exact Test with Bonferroni correction). By pairwise comparisons of the 70° × 3HIT datasets, we found that *w*^*1118*^ flies suffered a greater proportion of death by 0.5 h post-TBI than *y*^*1*^*w*^*1*^ flies (p = 0.001, α = 0.0125 by 2×2 Fisher’s Exact Test with Bonferroni correction). Alternately, *y*^*1*^*w*^*1*^ flies suffered greater death during the 2+ h to 8 h period than *w*^*1118*^ flies by comparison of 80° × 2HITs datasets (p = 0.004, α = 0.0125 by 2×2 Fisher’s Exact Test with Bonferroni correction).

### Time-Course of Motor Dysfunction

Flies are well-known to exhibit motor dysfunction following TBI (Katzenberger et al. 2013; Barekat et al. 2016; Anderson et al. 2018). However, the time-course of motor dysfunction across the primary and secondary injury periods, and for TBI of varying severities is not well-characterized. Therefore, we extended our characterization of outcomes across several time-points and the 70° × 3HITs, 80° × 2HITs, and 90° × 1HIT conditions via a negative geotaxis assay. We restricted our analysis to *w*^*1118*^ flies which have been best-characterized via this assay to date. We also opted to use a 6 h post-TBI time-point in place of the 8 h time-point used for the time-course of mortality to better capture outcomes during the middle of the peak of the secondary injury window.

We determined geotaxis performance by measuring the distance traveled by each fly 5 seconds and 10 seconds after startle, which were similar time-points to previous literature (Gargano et al. 2005; Linderman et al. 2012; Podratz et al. 2013; Anderson et al. 2018). All conditions showed improved scores at 10 s compared to 5 s (Fig. 5, p < 0.05, α = 0.5, Kruskal-Wallis with Dunn’s correction). In comparing conditions, we found that *w*^*1118*^ flies subjected to any of the three TBI conditions showed impaired geotaxis compared to uninjured flies at both the 0.5 h and 6 h post-TBI time-points, but not 2 h or 24 h post-TBI time-points, for each of the 5 s and 10 s measures (Fig. 5, see (*)). However, we observed no differences in geotaxis performance between any of the TBI conditions at any of the time-points tested. Comparisons within TBI conditions and across time-points showed a general trend for improved geotaxis scores at 2 h compared to 0.5 h post-TBI (Fig. 5A: see ‘a’ and ‘b’, Fig. 5B: see ‘a’, ‘b’, and ‘d’), followed by a second impairment in geotaxis for both the 70° × 3HITs and 80° × 2HITs datasets at 6 h post-TBI when assessed at 5 s post-startle (Fig. 5A, see ‘c’ and ‘d’).

**Figure 5:**
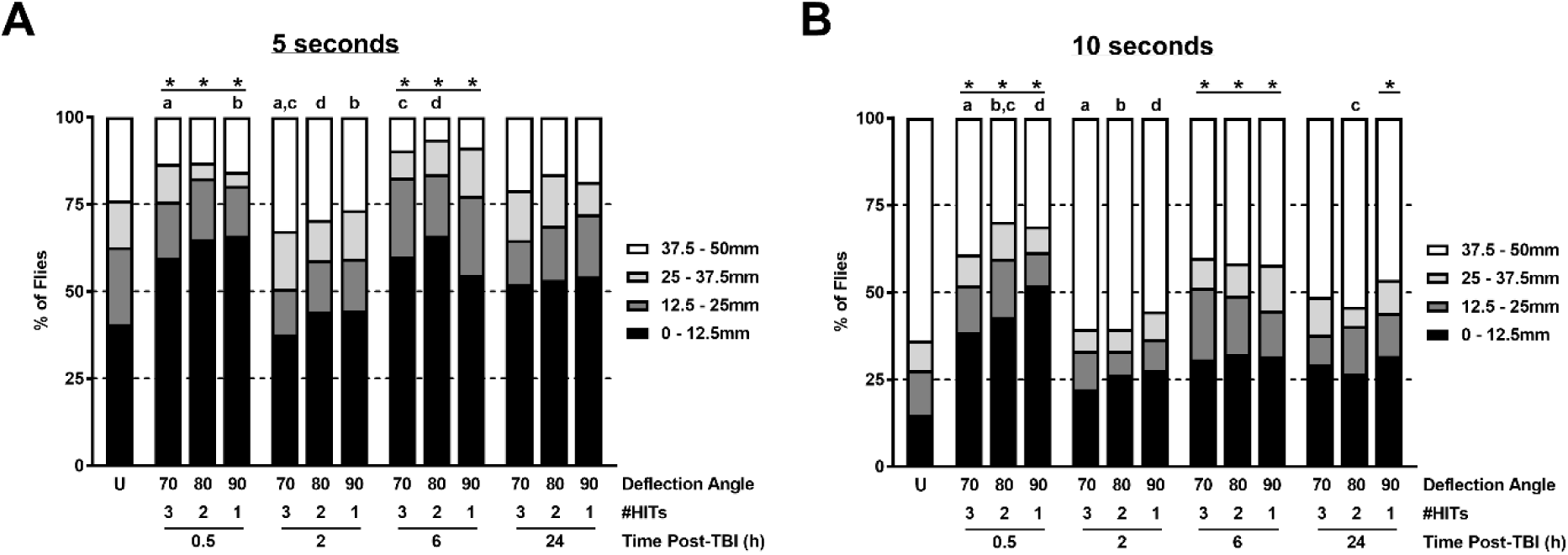
*w*^*1118*^ flies subjected to TBI exhibit time-dependent geotaxis impairment. Distances traveled by flies at 5 seconds (A) and 10 seconds (B) after startle were binned into 12.5mm quartiles for uninjured flies (U) and each TBI condition and assessed time post-TBI (h) indicated. Conditions that share a letter are statistically different, while (*) indicates a statistical difference between uninjured flies and the indicated condition (p ≤ 0.029, α = 0.05 by Kruskal-Wallis with Dunn’s correction). n ≥ 101 flies per condition.

## DISCUSSION

Fruit flies offer an accessible model to study TBI. Two models of conducting TBI studies in fruit flies are the high-impact trauma (HIT) method and the Bead Ruptor method (Katzenberger et al. 2013; Barekat et al. 2016). One advantage to the Bead Ruptor method is the ease of scaling the primary injuries (Barekat et al. 2016). We addressed this gap in methodology and expanded upon the original HIT method by adding selectable stopping points to reproducibly perform injury at four levels of injury severity. We then applied our expanded methodology to characterization of the time-course of death and locomotor dysfunction in the 24 h following TBI.

Several of our main findings are in agreement with the established TBI models. First, we found that increasing the injury number results in dose-dependent increases in mortality (Figs. 1 and 3) (Katzenberger et al. 2013; Barekat et al. 2016; Anderson et al. 2018). Second, we found that *y*^*1*^*w*^*1*^ flies suffer greater mortality than *w*^*1118*^ flies subjected to the same injuries, and we extended this finding to our mild and moderate TBI conditions (Figs. 1 and 2) (Katzenberger et al. 2013). Notably, our MI_24_ values at the standard protocol of 90° × 4HITs (Table 1, *w*^*1118*^: 53.3%; *y*^*1*^*w*^*1*^: 81.6%) were not identical to previous literature (*w*^*1118*^: ∼30%; *y*^*1*^*w*^*1*^: ∼50%) (Katzenberger et al. 2013; Anderson et al. 2018), likely due to lab specific differences such as variation in the force generated by the spring and/or the features of the collision surface. However, it is notable that we reproduced the data showing increased sensitivity of *y*^*1*^*w*^*1*^ flies (Katzenberger et al. 2013), thereby demonstrating the reliability of this TBI model and the penetrance of unknown genetic influences (Katzenberger et al. 2015). Third, we found that reducing the angle of deflection resulted in less severe primary injuries as indicated by decreased mortality, and extended this finding across the four levels of deflection tested (Fig. 2) (Anderson et al. 2018). Fourth, we found that TBI led to diminished locomotor ability that returned to normal levels by 24 h post-TBI (Fig. 5) (Katzenberger et al. 2013).

Several of our findings are also novel for this TBI system. First, we found a synergistic effect of additional HITs on mortality by our moderate TBI conditions and short inter-injury intervals (15 seconds) (Table 2). It was previously reported that dividing the MI_24_ by the number of HITs resulted in no differences when comparing across HIT number (Katzenberger et al. 2013). This result was used as evidence that the main factor influencing MI_24_ across multiple HITs was the likelihood of suffering a critical injury for each HIT, and that secondary injury mechanisms were negligible for injuries spaced closely together as the secondary injury window does not peak until 1-8 h after injury (Katzenberger et al. 2013, 2016). By contrast, we found differences when comparing median MI_24_/HIT values even at these close (15 second) inter-injury intervals (Figs. 1B, 1C, 3A-F). Additionally, we looked more closely at the pattern of MI_24_/HIT values across HIT number. If only primary injuries, and not secondary injuries or increased susceptibility to mortality due to preceding strikes, were responsible for observed MI_24_ values then the MI_24_/HIT values across HIT number should generate a zero slope line. At 90° neither *w*^*1118*^ nor *y*^*1*^*w*^*1*^ had significantly non-zero best-fit line slopes, consistent with properties of the primary injury being most responsible for MI_24_ at these severe injury levels and short inter-injury interval (Table 2).

By contrast, for our moderate severity injuries at 80° and 70°, both *w*^*1118*^ and *y*^*1*^*w*^*1*^ MI_24_/HIT data generated positive, significantly non-zero slopes, indicating a synergistic effect of HIT number on mortality. This result suggests that secondary injury mechanisms, or increased susceptibility to injury due to preceding injuries, contributed to MI_24_ (Table 2). A non-zero trend in MI_24_/HIT data was not observed for injuries at 60°, though it is possible that such a trend would be evident if injury number was further increased as the MI_24_ value increased noticeably between 3 and 4 HITs for both *w*^*1118*^ and *y*^*1*^*w*^*1*^ (Table 2).

A second novel finding was the pattern of death in the 24 h following TBI. First, we found a difference in the pattern of death when comparing 80° × 2HITs to 90° × 1HIT within both *w*^*1118*^ and *y*^*1*^*w*^*1*^ genotypes (Fig. 4). Interestingly, we observed an increased proportion of death in the late period from 8+ h to 24 h post-TBI in the 80° × 2HIT dataset for *y*^*1*^*w*^*1*^ flies (Fig. 4A), and a similar, nonsignificant trend for *w*^*1118*^ flies (Fig. 4B). This finding is consistent with our initial hypothesis that repetitive, moderate injury would lead to a greater proportion of death in the secondary injury period than a single, severe TBI event. However, by contrast to this finding, we found no differences when comparing the pattern of death via 70° × 3HITs to 90° × 1HIT (Fig. 4).

Why might 2HITs at the 80° deflection angle, but not 3HITs at 70°, cause a different pattern of death than 1HIT at 90°? One possibility is the presence of separate secondary injury mechanisms, one operating on a short time-scale (seconds) and the other the previously defined mechanism operating on a longer time-scale (hours) (Katzenberger et al. 2016). By this model, the administration of 2HITs at 80° causes significant secondary injuries on the longer time-scale and greater death after 8+ h compared to 1HIT at 90° (Fig. 4B). By contrast, 70° × 3HITs and the synergistic secondary effects on the short time-scale exceeds a critical threshold and causes early death before the classic secondary injury period. Such a two-stage secondary injury model may also be applicable to differences between genotypes. By our data, the most common period of death for *w*^*1118*^ flies regardless of TBI condition was within 0.5 h of injury, while for *y*^*1*^*w*^*1*^ flies it was during the typical secondary injury period from 2+ h to 8 h for 70° × 3HITs and 90° × 1HIT datasets and 8+ h to 24 h for the 80° × 2HITs dataset. This data suggests that *w*^*1118*^ flies are more susceptible to fast secondary injury mechanisms (seconds), while *y*^*1*^*w*^*1*^ flies are more susceptible to the later secondary injury mechanisms (hours). Consistent with this, administration of 70° × 3HITs to each genotype revealed a significant increase in the proportion of death by the 0.5 h time-period for *w*^*1118*^ as compared to the proportion for *y*^*1*^*w*^*1*^ flies (Fig. 4), consistent with a threshold sensitive to fast secondary injury mechanisms in the genotype more sensitive to these changes. By contrast, the 80° × 2HIT condition which may not exceed the early threshold, caused an increase in death during the later, classic secondary injury period from 2+ h to 8 h in *y*^*1*^*w*^*1*^ flies more susceptible to these mechanisms.

Our final novel finding was the more detailed pattern of locomotor dysfunction in the 24 h following TBI and for isolated vs repetitive TBI conditions. We found that flies showed early locomotor dysfunction at 0.5 h compared to uninjured flies for all three TBI conditions tested, but found no differences between TBI conditions at this early time-point (Fig. 5). We then saw a trend for improved geotaxis scores that were no different than controls by the early stages of the classic secondary injury period at 2 h post-TBI and during the delayed secondary injury period 24 h post-TBI, consistent with previous results for 1-2HITs at 90° (Katzenberger et al. 2013) (Fig. 5). More interesting and novel were the impaired geotaxis scores for injured flies at 6 h post-TBI, a time corresponding to the middle of the secondary injury period. However, diminished geotaxis scores at 6 h post-TBI were only different than 2 h scores for the repetitive TBI conditions of 70° × 3HITs and 80° × 2HITs (Fig. 5A). Thus, all TBI conditions led to early geotaxis impairment and a second impairment during the secondary injury window at 6 h post-TBI, while the secondary period difference at 6 h post-TBI was most noted in the repetitive TBI conditions.

What are the mechanisms by which closely spaced, mild or moderate injuries synergistically affect TBI outcomes? Secondary mechanisms might include autophagy-related pathways and stress granule formation (Anderson et al. 2018). In fly larvae, stress granules were not apparent after single TBI events at 60°, minimally increased after 4HITs, and substantially increased after 8HITs in an apparently synergistic fashion (Anderson et al. 2018). Alternatively, glutamate release and elevated extracellular potassium are observed immediately or within minutes of TBI (Faden et al. 1989; Katayama et al. 1990). Moreover, extracellular potassium scaled with injury severity until plateauing for severe injuries, and changes in extracellular potassium were blocked by addition of tetrodotoxin for moderate but not severe injuries (Katayama et al. 1990). Thus, dysregulation of neuronal excitability and extracellular potassium operate on short time-scales and are responsive to injury severity, which are compatible with our observed synergistic effects for injuries at short inter-injury intervals and for moderate, but not severe TBI. Downstream consequences of misregulated neurotransmission and extracellular potassium are varied, but may include changes in oxidative stress and inflammation (Guerriero et al. 2015; Fehily and Fitzgerald 2017; Khatri et al. 2018).

Our evidence for synergistic effects of mild to moderate TBI and short inter-injury intervals is of consequence to mammals. In mammals the secondary injury window is typically reported as within days post-injury (Laurer et al. 2001; Longhi et al. 2005; Friess et al. 2009; Meehan et al. 2012; Huang et al. 2013; Bolton and Saatman 2014; Weil et al. 2014; Bolton Hall et al. 2016). However, the number of sub-concussive events suffered by individuals across a short time-scale, a single American football game, correlated with short-term blood-brain-barrier damage (Marchi et al. 2013). Thus, mild TBI events suffered in number across a short time-scale may be an important factor to consider for brain health, especially considering the large number (> 1,000) of sub-concussive injuries suffered during football participation across a season of play (Crisco et al. 2010).

The exact secondary mechanisms underlying the fast synergistic effects we observed are thus far unknown. Moreover, we do not know if synergistic effects at mild to moderate TBI conditions in our model drive other TBI-related consequences observed in flies such as changes in lifespan and inflammation (Katzenberger et al. 2013, 2015, 2016; Barekat et al. 2016; Anderson et al. 2018). However, our fly model offers an unparalleled platform for rapidly, and systematically, testing candidate factors or pathways for their involvement in TBI outcomes across injury severities, number, and inter-injury interval. Recognition and elucidation of cellular and molecular differences in response to mild vs moderate, single vs multiple TBI events, and across varying time-scales will be important in determining optimal disease-intervention strategies. Our extended model will be central to these efforts going forward.

## Acknowledgements

The authors thank members of the Human Biology Department at the University of Wisconsin – Green Bay for careful reading of the manuscript and suggested revisions. We also thank Dr. C. Andrew Frank, Dr. Tina Tootle, Dr. Atulya Iyengar, and Dr. Javier Gomez for reading the manuscript and offering revisions, and for their support in answering many questions about lab set-up and publication. We also thank Mark Damie and Joe Schoenebeck of the University of Wisconsin – Green Bay for assistance in lab set-up.

## Declarations of Interests

The authors have no conflicts of interest.

## Funding

This work was supported by University of Wisconsin – Green Bay Start-Up Funds, University of Wisconsin – Green Bay Summer Scholar Grant, and Medical College of Wisconsin Professional Development Funds.

## Notes

#### Summary of Updates

Characterization of the time-course of death (Fig 4) and locomotor dysfunction (Fig 5) in the 24 h following injury was added. Textual edits.

